# Converging Evidence from Electrocorticography and BOLD fMRI for a Sharp Functional Boundary in Superior Temporal Gyrus Related to Multisensory Speech Processing

**DOI:** 10.1101/272823

**Authors:** Muge Ozker, Michael S. Beauchamp

## Abstract

Although humans can understand speech using the auditory modality alone, in noisy environments visual speech information from the talker’s mouth can rescue otherwise unintelligible auditory speech. To investigate the neural substrates of multisensory speech perception, we recorded neural activity from the human superior temporal gyrus using two very different techniques: either directly, using surface electrodes implanted in five participants with epilepsy (electrocorticography, ECOG), or indirectly, using blood oxygen level dependent functional magnetic resonance imaging (BOLD fMRI) in six healthy control fMRI participants. Both ECOG and fMRI participants viewed the same clear and noisy audiovisual speech stimuli and performed the same speech recognition task. Both techniques demonstrated a sharp functional boundary in the STG, which corresponded to an anatomical boundary defined by the posterior edge of Heschl’s gyrus. On the anterior side of the boundary, cortex responded more strongly to clear audiovisual speech than to noisy audiovisual speech, suggesting that anterior STG is primarily involved in processing unisensory auditory speech. On the posterior side of the boundary, cortex preferred noisy audiovisual speech or showed no preference and showed robust responses to auditory-only and visual-only speech, suggesting that posterior STG is specialized for processing multisensory audiovisual speech. For both ECOG and fMRI, the transition between the functionally distinct regions happened within 10 mm of anterior-to-posterior distance along the STG. We relate this boundary to the multisensory neural code underlying speech perception and propose that it represents an important functional division within the human speech perception network.

## Introduction

The human ability to understand speech is one of our most important cognitive abilities. While speech can be understood using the auditory modality alone, vision provides important additional cues about speech. In particular, the mouth movements made by the talker can compensate for degraded or noisy auditory speech (Bernstein et al 2004, Ross et al 2007, Sumby & Pollack 1954). While it has been known since Wernicke that posterior lateral temporal cortex is important for language comprehension, the advent of blood-oxygen level dependent functional magnetic resonance imaging (BOLD fMRI) led to important advances, such as the discovery that multiple regions in temporal cortex are selective for human voices (Belin et al 2000). However, BOLD fMRI suffers from a major limitation, in that it is a slow and indirect measure of neural function. Spoken speech contains 5 or more syllables per second, requiring the neural processes that decode each syllable to be completed in less than two hundred milliseconds. In contrast, the sluggish hemodynamic response that underlies BOLD fMRI does not peak until several seconds after the neural activity that prompted it.

This drawback underscores the importance of complementing fMRI with other techniques that directly measure neural activity. The non-invasive techniques of EEG and MEG have led to a better understanding of the temporal dynamics of speech perception (Crosse et al 2016, Salmelin 2007, Shahin et al 2012, Sohoglu & Davis 2016). Recently, there has also been tremendous interest in electrocorticography (ECOG), a technique in which electrodes are implanted in the brains of patients with medically intractable epilepsy. Compared with EEG and MEG, ECOG allows localization of activity to the small population of neurons nearest each electrode, leading to the discovery of selective responses in the superior temporal gyrus (STG) for various speech features, including categorical representations of speech (Chang et al 2010) phonetic features (Mesgarani et al 2014) and prosody (Tang et al 2017).

While the broad outlines of the organization of visual cortex are well-established (Grill-Spector & Malach 2004), the layout of auditory cortex is less well known. Early areas of auditory cortex centered on Heschl’s gyrus contains maps of auditory frequency and spectral temporal modulation (Moerel et al 2014). In contrast, within auditory association cortex in the STG, organization by auditory features is weaker, and the location and number of different functional areas is a matter of controversy (Leaver & Rauschecker 2016). Recently, we used ECOG to document a double dissociation between anterior and posterior regions of the STG (Ozker et al 2017). Both regions showed strong responses to audiovisual speech, but the anterior area strongly preferred speech in which the auditory component was clear while the posterior area preferred speech in which the auditory component was noisy or showed no preference. There was a sharp anatomical boundary, defined by the posterior edge of Heschl’s gyrus, between the two areas. All electrodes anterior to the boundary preferred clear speech, and no electrodes posterior to the boundary did. These results were interpreted in the conceptual framework of multisensory integration. Auditory association areas in anterior STG respond strongly to clear auditory speech but show a reduced response because of the reduced information available in noisy auditory speech, paralleling the reduction in speech intelligibility. Multisensory areas in posterior STG area able use the visual speech information to compensate for the noisy auditory speech, restoring intelligibility. However, this demands recruitment of additional neuronal resources, leading to an increased response during noisy audiovisual speech perception.

While there have been numerous previous fMRI studies of noisy and clear audiovisual speech, *e.g.* (Bishop & Miller 2009, Callan et al 2003, Lee & Noppeney 2011, McGettigan et al 2012, Sekiyama et al 2003, Stevenson & James 2009) none described a sharp boundary in the response patterns to clear and noisy speech within the STG. BOLD fMRI has the spatial resolution necessary to detect fine-scale cortical boundaries, such as between neighboring ocular dominance columns (Cheng et al 2001), ruling out sensitivity of the technique itself as an explanation. Instead, we considered two other possibilities. One possible explanation is that the analysis or reporting strategies used in previous fMRI studies (such as group averaging or reporting only activation peaks) could have obscured a sharp functional boundary present in the fMRI data. A second, more worrisome, explanation is that the sharp boundary observed with ECOG reflects anomalous brain organization in the ECOG participants. Brain reorganization due to repeated seizures could have resulted in different STG functional properties in epileptic patients compared with healthy controls (Janszky et al 2003, Kramer & Cash 2012).

To distinguish these possibilities, we collected BOLD fMRI data from healthy controls viewing the same clear and noisy audiovisual speech stimuli viewed by the ECOG patients. Both fMRI and ECOG participants performed the same speech identification task. The BOLD fMRI data was analyzed without any spatial blurring or group averaging to ensure that these would not obscure areal boundaries within the STG. fMRI allows sampling of the entire brain volume, instead of the limited coverage obtained with ECOG electrodes. We used this ability to examine the preference for clear or noisy speech across the entire length of the STG.

## Methods

All participants provided written informed consent and underwent experimental procedures approved by the Baylor College of Medicine (BCM) Institutional Review Board. ECOG data was collected from five participants with refractory epilepsy (3 women, mean age 31 years) and fMRI data was collected from six participants recruited from the BCM community (3 women, mean age 25 years).

### Stimulus and Task

For the main experiment, identical stimuli were used for the ECOG and fMRI participants. The stimuli consisted of audiovisual recordings of a female talker from the Hoosier Audiovisual Multi-Talker Database speaking single words (“rain” or “rock”) in which the auditory component of was either unaltered (auditory-clear) or replaced with speech-specific noise that matched the spectrotemporal power distribution of the original auditory speech (auditory-noisy). A parallel manipulation was performed on the visual component of the speech by replacing the original video with a highly blurred version, resulting in four conditions (auditory-clear + visual-clear; auditory-clear + visual-blurred; auditory-noisy + visual-clear; auditory-noisy + visual-blurred). For the ECOG participants, from 32 to 56 repetitions of each condition were presented in random order. For the fMRI participants, 60 repetitions of each condition were presented in random order.

Following each stimulus presentation, participants performed a two-alternative forced choice on the identity of the presented word. Accuracy was at ceiling for the auditory-clear conditions (auditory-clear with visual-clear: 99% for fMRI participants, 99% for ECOG participants; with visual-noisy: 99% for fMRI, 98% for ECOG) and lower for auditory-noisy conditions (with visual-clear: 91% for fMRI, 81% for ECOG; with visual-noisy: 75% for fMRI, 63% for ECOG).

fMRI participants also passively viewed standard localizer stimuli consisting of auditory-only, visual-only and audiovisual version of short stories (Aesop’s fables) (Nath & Beauchamp 2012, Nath et al 2011).

### Definition of Anterior and Posterior Superior Temporal Gyrus (STG)

Cortical surface models were constructed from the high-resolution T1-weighted anatomical MRI scans of ECOG and fMRI participants using FreeSurfer (Fischl et al 2001). For ECOG participants, the post-implantation computed tomography (CT) scan showing the electrode locations was registered to the anatomical MRI to ensure accurate electrode localization.

Two atlases were used to parcellate the STG. The Destrieux atlas defines the entire STG using the “G_temp_sup-Lateral” label (lateral aspect of the STG) (Destrieux et al 2010). The Desikan-Killiany atlas (Desikan et al 2006) applies a single “Superior Temporal” label to both the STG and the STS with an additional “Banks Superior Temporal” label for the posterior portion of the STS, with an anterior border defined by the posterior-most point of Heschl’s gyrus. We cleaved the Destrieux STG into an anterior portion and a posterior portion using the Heschl’s gyrus landmark defined by the Desikan-Killiany atlas (boundary shown as black dashed lines in Figure 1). The posterior STG is continuous with the supramarginal gyrus. Since the two atlases vary in their handling of this boundary, we manually defined the posterior boundary of the STG as being just past the location where the gyrus begins its sharp turn upward into parietal lobe. All analyses were done only within single participants without any normalization or spatial blurring. In order to report the location of the anterior-posterior boundary in standard space, individual MRIs were aligned to the N27 brain (Holmes et al 1998).

**Figure 1:**
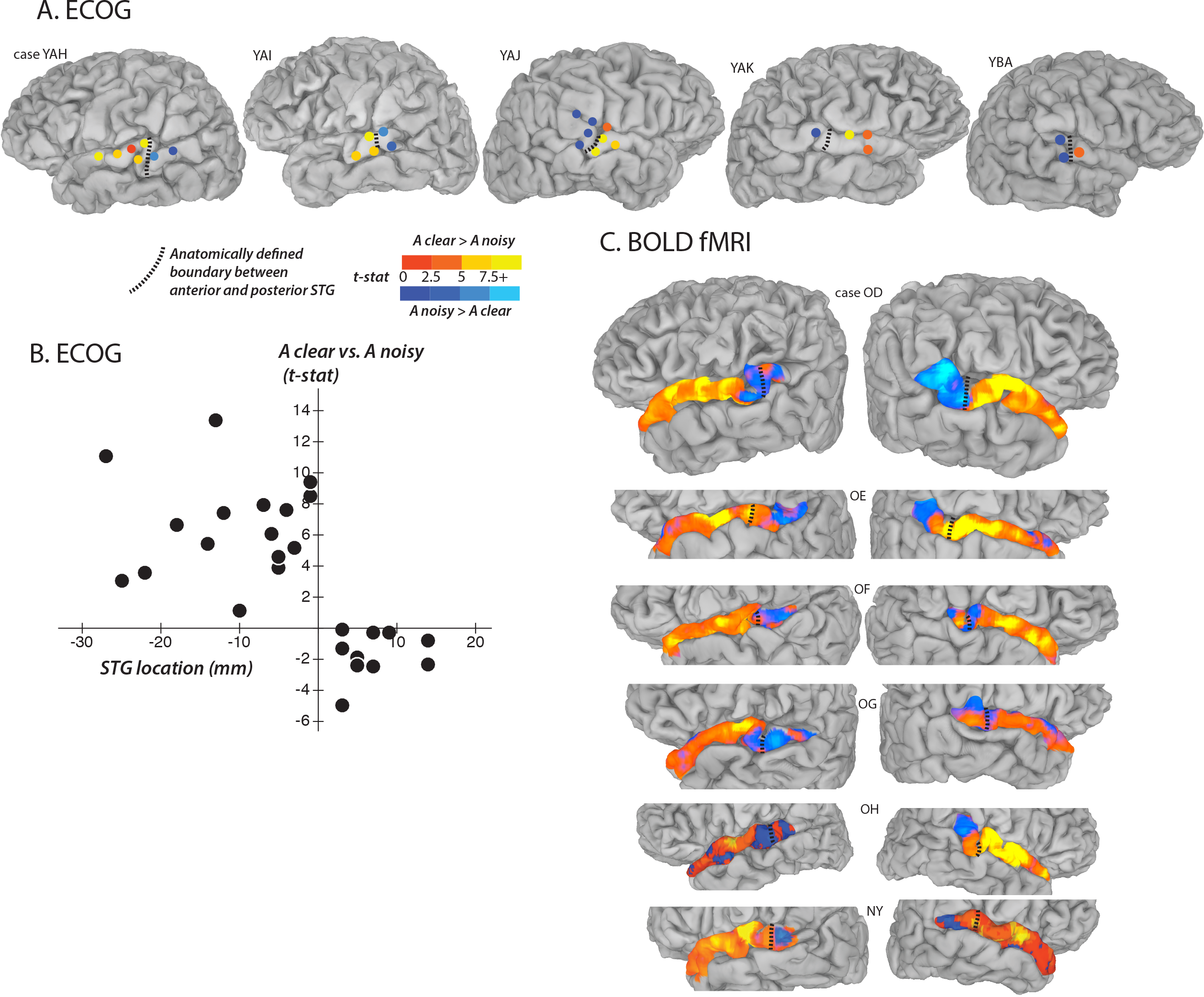
Converging evidence from fMRI and ECOG for a functional boundary in posterior STG. A. Cortical surface models of five hemispheres from five ECOG participants (case letter codes indicate anonymized participant IDs). Colored circles show locations of subdural electrodes on the STG showing a significant response to audiovisual speech. Warm electrode colors indicate greater response to audiovisual speech with a clear auditory component. Cool electrode colors indicate greater response to speech with a noisy auditory component. Dashed black line shows the location of the anatomical border between anterior STG and posterior STG defined by the Desikan-Killiany atlas (Desikan et al 2006). B. The preference of each electrode for clear or noisy speech (y-axis) plotted against its location relative to the anterior-posterior STG border (x-axis) with one symbol per electrode. Negative values on the x-axis are more anterior, positive values are more posterior. C. Cortical surface models of twelve hemispheres from six fMRI participants. Surface nodes on the STG are colored by their preference for clear or noisy audiovisual speech (same color scale for A and C).

### ECOG Experimental Design and Data Analysis

Experiments were conducted in the epilepsy monitoring unit of Baylor St. Luke’s Medical Center. Patients rested comfortably in their hospital beds while viewing stimuli presented on an LCD monitor mounted on a table and positioned at 57 cm distance from the participant. While the participants viewed stimulus movies, a 128-channel Cerebus amplifier (Blackrock Microsystems, Salt Lake City, UT) recorded from subdural electrodes that consisted of platinum alloy discs (diameter 2.3 mm) embedded in a flexible silicon sheet with inter-electrode distance of 10 mm. An inactive intracranial electrode implanted facing the skull was used as a reference for recording. Signals were amplified, filtered, and digitized at 2 kHz. Offline, common average referencing was used to remove artifacts, and the data was epoched according to stimulus timing. Line noise was removed and spectral decomposition was performed using multitapers. The measure of neural activity was the broad-band high-gamma response (70 - 110 Hz) measured as the percent change relative to a pre-stimulus baseline window (500 to 100 ms before auditory stimulus onset). The high broadband response was used as it is the ECOG signal most closely associated with the rate of action potentials and the BOLD fMRI response (reviewed in Ojemann et al 2013, Ray & Maunsell 2011). Across patients, a total of 527 intracranial electrodes were recorded from. Of these, 55 were located on the STG. 27 of these showed a minimal level of stimulus-related activity, defined as significant high-gamma responses to audiovisual speech compared with prestimulus baseline (*p* < 10^−3^, equivalent to ~40% increase in stimulus power from baseline) and were included in the analysis.

### fMRI Experimental Design and Data Analysis

Experiments were conducted in the Core for Advanced MRI (CAMRI) at Baylor College of Medicine using a 3 Tesla Siemens Trio MR scanner equipped with a 32-channel head gradient coil. BOLD fMRI data was collected using a multislice echo planar imaging sequence (Setsompop et al. 2012) with TR = 1500 ms, TE = 30 ms, flip angle = 72°, in-plane resolution of 2 × 2 mm, 69 2-mm axial slices, multiband factor: 3, GRAPPA factor: 2. fMRI data was analyzed using the afni_proc.py pipeline (Cox 1996). Data was time shifted to account for different acquisition times for different slices; aligned to the first functional volume which was in turn aligned with the high-resolution anatomical; and rescaled so that each voxel had a mean of 100. No blurring or spatial normalization of any sort was applied to the EPI data.

Data for the main fMRI experiment was collected in five runs, each with 160 brain volumes (4 minutes duration). Each run contained forty-eight 3-second trials, twelve for each stimulus condition, for a total of sixty repetitions of each condition. A rapid event-related design was used with fixation baseline occupying the remaining 96 seconds of each run, optimized with the scheduling algorithm optseq2 (https://surfer.nmr.mgh.harvard.edu/optseq) (Dale et al 1999).

Data for the localizer fMRI experiment was collected in two runs, each with 180 brain volumes (4:30 duration); for one participant, only one run was collected. Each run contained nine blocks (20-seconds of stimulus, 10 seconds of fixation baseline) consisting of three blocks each (in random order) of auditory-only, visual-only, and audiovisual speech recorded by a female talker (Nath & Beauchamp 2012, Nath et al 2011).

A generalized linear model was used to model the fMRI time series independently for each voxel using the 3dDeconvolve function in AFNI. The model contained 10 regressors: 6 regressors of no interest generated by the motion correction process and 4 regressors of interest (one for each stimulus condition) using an exponential hemodynamic response function generated with the 3dDeconvolve option “BLOCK(2,1)”. A general linear test with the values of “+1 +1 −1 −1” was used to find the *t*-statistic for the contrast between the two conditions with clear auditory speech and the two conditions with noisy auditory speech (data in Figures 1 and 2). This contrast between auditory-clear and auditory-noisy was the main dependent measure in the analysis.

**Figure 2.**
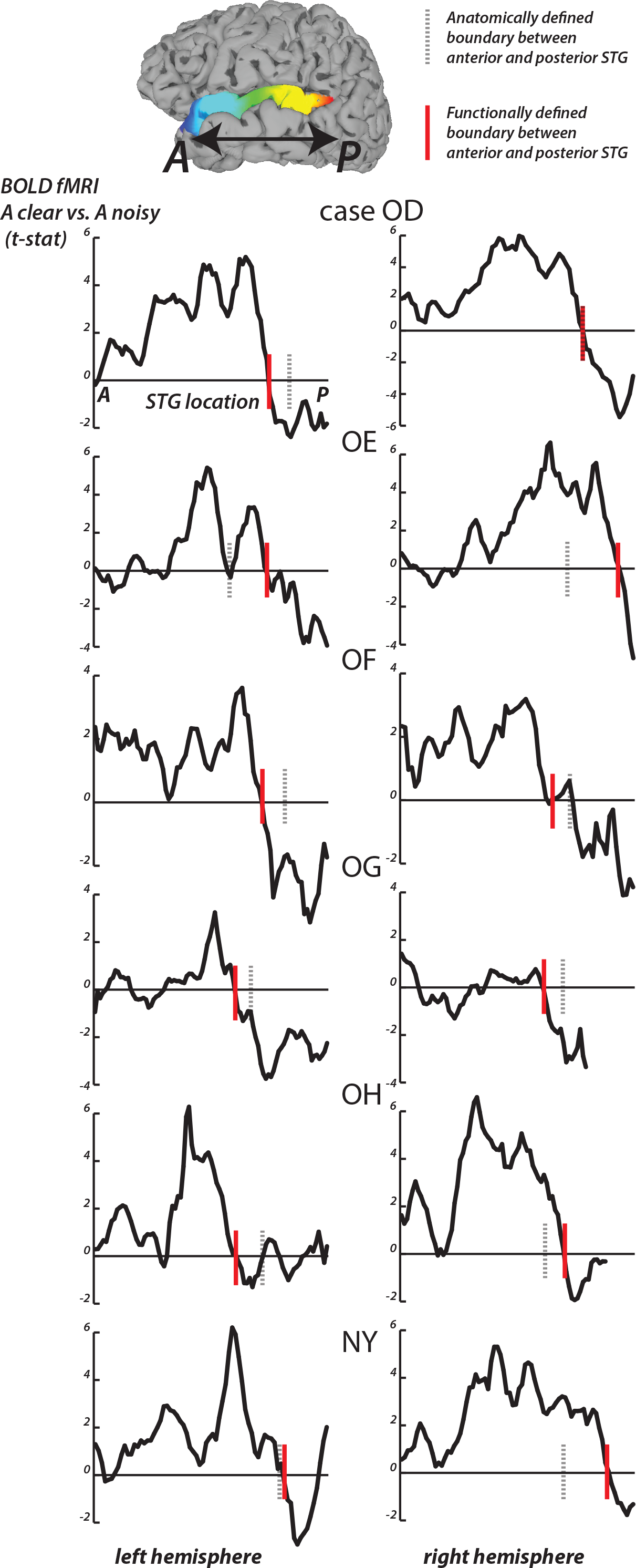
fMRI responses along the length of the STG. For each fMRI participant, the STG was parcellated into 1 mm bins from the most anterior point (A) to the most posterior point (P). For all surface nodes in each bin, the average value of the clear *vs*. noisy *t*-statistic was averaged. In each plot, the y-axis is the average *t*-statistic and the x-axis is the location along the STG from A to P. The left column shows plots from the left hemispheres, the right column shows plots for the right hemispheres; case letter codes indicate anonymized participant IDs. In each hemisphere, the location of the functional boundary between anterior STG and posterior STG was defined as the first zero-crossing of this curve in the posterior third of the STG (red vertical lines). In each hemisphere, the anatomical boundary between anterior and posterior STG was defined by the posterior margin of Heschl’s gyrus (gray dashed vertical lines). For case OD right hemisphere, the two boundaries overlap.

For the STG length analysis (Figure 2), unthresholded fMRI data in the form of the clear *vs*. noisy *t*-statistic was mapped to the cortical surface using the AFNI function 3dVol2Surf. The options “−ave −f_steps 15” were used, resulting in a line between each node on the pial surface and the corresponding node on the smoothed white matter surface being subdivided into 15 equal segments, with the fMRI voxel values at each segment sample and averaged. The entire STG was divided into 1 mm bins, from anterior to posterior, and the *t*-statistic at all nodes within each bin was averaged. For each hemisphere, a functional boundary was defined as the bin containing the first zero-crossing of the *t-*statistic (moving in an anterior-to-posterior direction) in the posterior third of the STG.

To estimate the shape of the hemodynamic response function without assumptions (data in Figures 3, 4, 5), a second model was constructed that used tent functions to estimate the amplitude of the response independently at each time point of the BOLD response. For the main fMRI experiment, the response window spanned 0 to 15 seconds after stimulus onset (11 time points at a TR of 1.5 seconds) using the 3dDeconvolve option “TENTzero(0,15,11)”, resulting in a model with 50 regressors (6 motion regressors and 44 regressors of interest). For the localizer fMRI experiment, the response window spanned 0 to 30 seconds after stimulus onset (21 time points) using “TENTzero(0,30,21)” for each of the three block types. The beta-weights at the 4.5 second, 6 second, and 7.5 second time points were averaged to create an estimate of response amplitude.

**Figure 3.**
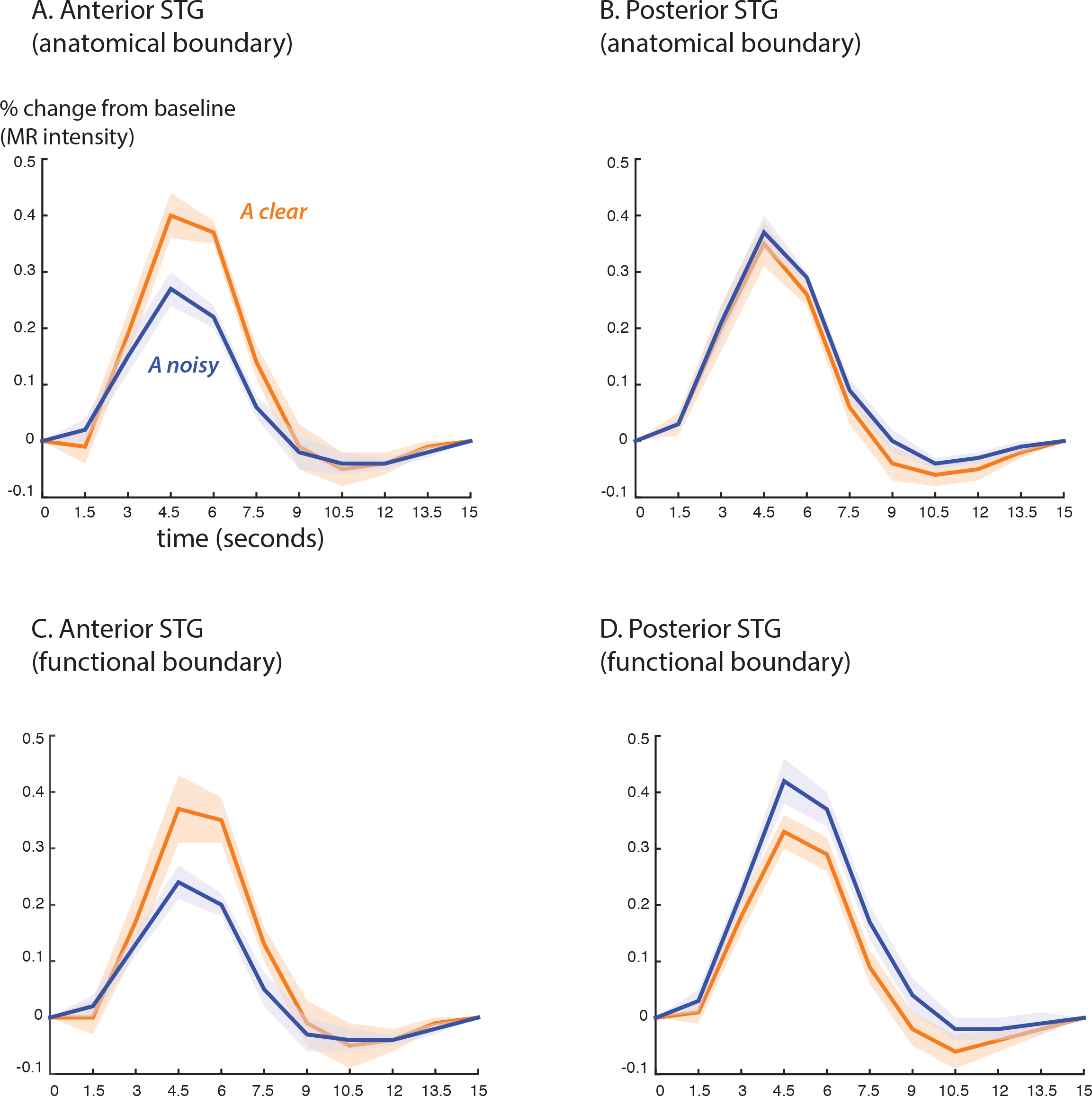
BOLD fMRI Responses to clear and noisy audiovisual speech in anterior and posterior STG. A. The average fMRI response for anterior STG was created by selecting all voxels in each hemisphere that were located from 0 mm to 30 mm anterior to the anatomical boundary defined by Heschl’s gyrus and that showed a significant response to any stimulus. Responses shown for audiovisual speech with a clear auditory component (blue) and a noisy auditory component (orange). Lines show mean, shaded regions show SEM across participants. B. The average fMRI response for posterior STG was created by selecting all responsive voxels in each hemisphere that were located from 0 mm to 15 mm posterior to the anatomical boundary defined by Heschl’s gyrus. C. Average BOLD fMRI response in the anterior STG, defined as all responsive voxels located from 0 mm to 30 mm anterior to the functional boundaries defined as shown in Figure 2. D. BOLD fMRI response in the posterior STG, defined as all responsive voxels located from 0 mm to 15 mm posterior to the functional boundary.

**Figure 4.**
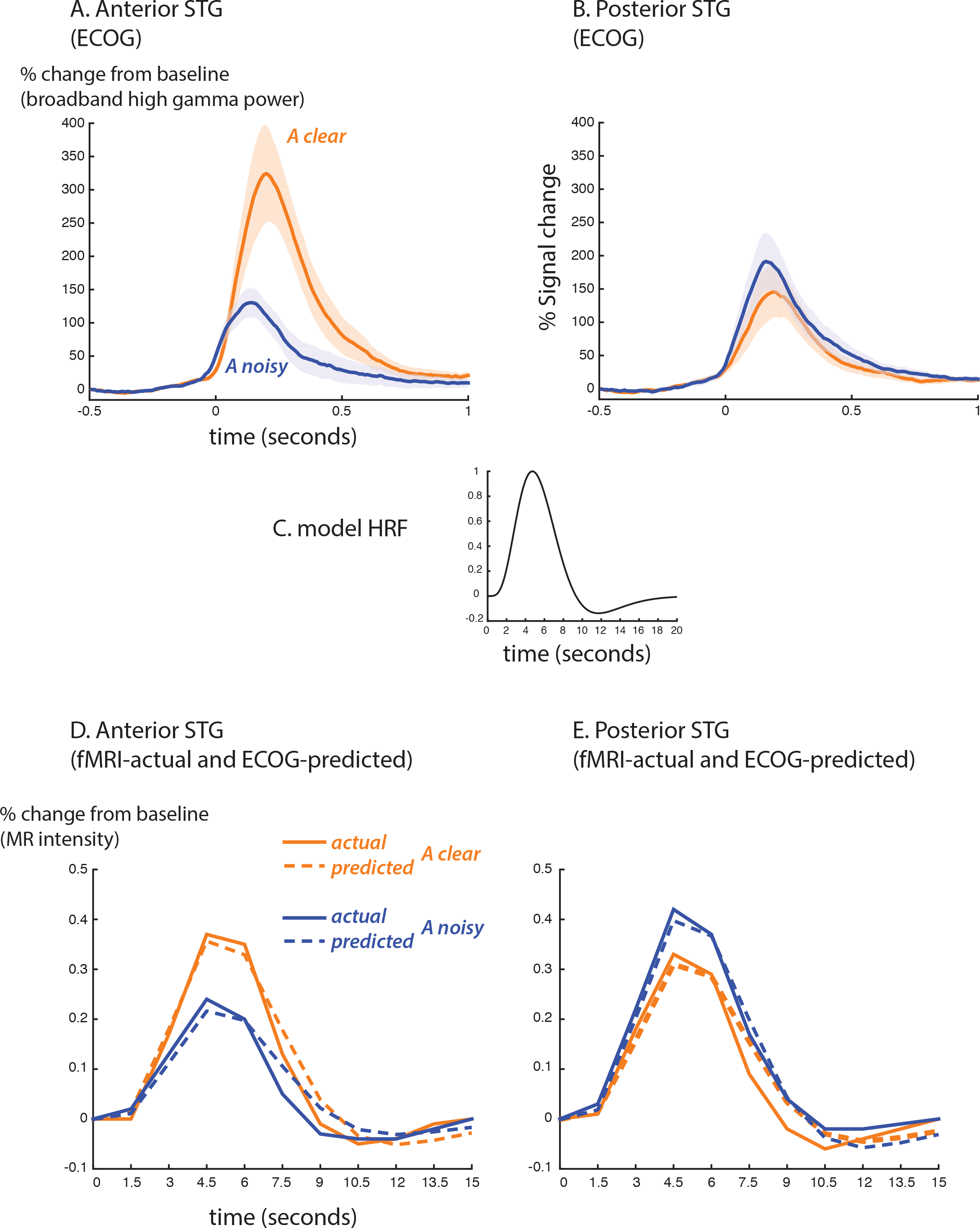
Comparison of ECOG and BOLD fMRI responses. A. The broadband high-gamma power (70 - 110 Hz) in the ECOG response plotted as the increase relative to prestimulus baseline (−500 to −100 ms) for audiovisual speech with a clear auditory component (blue line) and a noisy auditory component (orange line). Grand mean across all anterior STG electrodes in all participants (shaded region shows SEM). B. Grand mean response to clear and noisy speech across all posterior STG electrodes. C. A model hemodynamic response function (HRF) used to create the shape of the predicted fMRI response by convolving with the ECOG response. D. Predicted fMRI responses for anterior STG (dotted lines) compared with the actual fMRI responses from Figure 3C (solid lines). The predicted response was created by convolution with the model HRF and fitting a scale factor to determine the amplitude. A separate scale factor was used for each condition. E. Predicted fMRI responses for posterior STG (dotted lines) compared against the actual fMRI responses from Figure 3D (solid lines).

**Figure 5.**
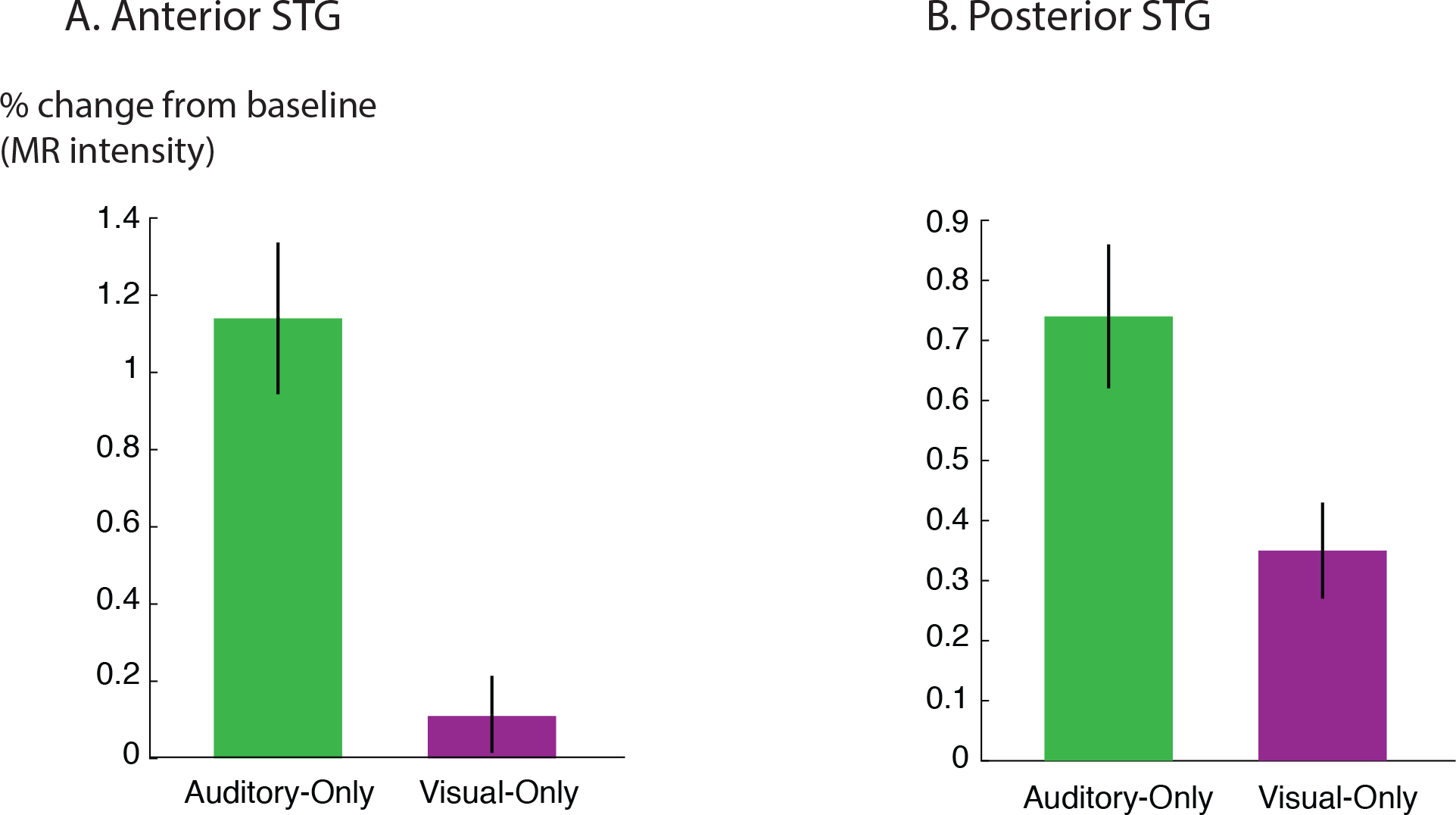
STG responses to unisensory auditory and visual speech. A. The response of anterior STG to unisensory auditory and visual speech in the fMRI localizer experiment. B. The response of posterior STG to unisensory auditory and visual speech in the fMRI localizer experiment.

To estimate the average BOLD fMRI hemodynamic response function, anterior and posterior STG ROIs were created in each participant using as a boundary either the anatomical Heschl’s gyrus boundary or the functional boundary defined by the STG length analysis. The anterior STG ROI contained all voxels from 0 to 30 mm anterior to the boundary and the posterior STG ROI contained all voxels from 0 to 15 mm posterior to the boundary. These values were chosen for consistency with the ECOG electrode locations, which ranged from 30 mm anterior to the anatomical/functional boundary to 15 mm posterior to it (Figure 1B). For correspondence with the ECOG electrode selection criteria (in which only electrodes that showed some response were included in the analysis) only voxels with an omnibus Full-F statistic of F > 5 (q < 10^−6^) were included in the ROIs.

To directly compare the BOLD fMRI with the ECOG responses, the ECOG response were convolved with a double-gamma hemodynamic response function with peak time = 6 seconds, undershoot time = 10 seconds and response-to-undershoot ratio = 4 (Lindquist et al 2009). The only free parameter was a scale parameter that matched the amplitude of the predicted and actual fMRI responses; ascale parameters that minimized the difference between the predicted and actual fMRI responses for each of the four curves were found using the Matlab function *fminbnd* (Figure 4).

### Linear Mixed-effects Models

Linear mixed-effect models were constructed using *R* with the lme4 package to understand the relationship between the different experimental variables. The dependent measure was the % signal change from baseline. The fixed factors were location (anterior *vs*. posterior STG), the presence or absence of auditory noise (auditory-clear *vs*. auditory-noisy) and the presence or absence of visual noise (visual-clear *vs*. visual-noisy). For each fixed factor, the LME estimated the significance of the effect and the magnitude of the effect relative to a baseline condition, which was always the response to auditory-clear, visual-clear speech in anterior STG.

### Results

Electrodes implanted on the posterior portion of the STG responded robustly to audiovisual speech. However, within the posterior STG we observed a striking dissociation between more anterior electrodes, which preferred audiovisual speech with a clear auditory component, and more posterior electrodes, which preferred speech with a noisy auditory component or showed no preference (Figure 1A). The posterior-most point of Heschl’s gyrus has been proposed as a boundary dividing the STG/STS into anterior and posterior sections (Desikan et al 2006, Ozker et al 2017). As shown in Figure 1B, adapted from Figure 2G of (Ozker et al 2017), all electrodes anterior to the boundary preferred clear speech while none of the electrodes posterior to the boundary did. The difference in response patterns between anterior and posterior electrodes was very large, even between electrodes that were only 10 mm apart, the closest possible distance in our recording array. For instance, in one participant the response to clear speech of an anterior electrode was double its response to noisy speech (138% ± 13% *vs*. 49% ± 5%, mean across trials ± SEM; unpaired t-test: t_109_ = +6.2, p = 10^−8^) while the adjacent electrode, located 10 mm posterior across the boundary, responded half as much to clear speech as noisy speech (38% ± 5% *vs*. 89% ± 9%, t_109_ = −4.5, p = 10^−5^).

To determine if a similar boundary could be observed with fMRI, we scanned participants viewing the same stimuli as the ECOG participants and mapped the unthresholded statistical contrast of clear *vs*. noisy speech to a cortical surface model of each hemisphere (Figure 1C). The most posterior portion of the STG preferred noisy speech or showed no preference while more anterior regions preferred clear speech.

To quantify the location of the boundary between anterior and posterior STG, the preference for clear *vs*. noisy audiovisual speech in unthresholded fMRI data was plotted in 1 mm bins for the entire anterior-to-posterior extent of the STG (Figure 2). In the posterior STG of each hemisphere, there was a sign change in the *t*-statistic of the clear *vs*. noisy contrast (red lines in Figure 2). This sign change was used to define a *functional* A-P boundary in the STG. The posterior margin of Heschl’s gyrus was used to define an *anatomical* A-P boundary in the STG (black lines in Figure 2).

The mean anterior-to-posterior location of the fMRI-defined functional boundary in standard space was y = −28 mm (+−9 mm SD). The mean standard space location of the atlas-defined anatomical boundary in these participants was y = −30 mm (+−5 mm SD).

In some cases, the boundaries aligned remarkably well (*e.g*. inter-boundary distance of 1 mm, case OD right hemisphere) while in others they were farther apart (*e.g*. distance of 20 mm, case OE right hemisphere). There was no consistent anterior-to-posterior difference between the anatomical and functional boundaries, resulting in a small mean distance between them (y = −28 *vs*. −30, *t*_11_ = 0.2, *p* = 0.8).

The location of the anatomical and functional boundaries in the fMRI participants were similar to that of the anatomical boundary in the ECOG participants, located at y = −27 mm (+−2 mm SD); the 1 cm spacing of the ECOG electrodes did not allow a separate estimate of the functional boundary.

As in the ECOG data, the fMRI transition between clear and noisy speech preferring cortex happened over a short cortical distance. For instance, in participant OD’s left hemisphere, the *t*-statistic of the clear *vs*. noisy contrast changed from *t* = +5 to *t* = −2 within 10 mm.

Next, we examined the fMRI response profiles on either side of the anatomical and functional boundaries (Figure 3). We classified the STG from 0 to 30 mm anterior to each boundary as “anterior” and the STG from 0 to 15 mm posterior to each boundary as “posterior”. These values were chosen for consistency with the ECOG electrode locations, which ranged from 30 mm anterior to the boundary to 15 mm posterior to it.

Both the anatomical and functionally-defined STG ROIs showed the characteristic BOLD response, with a positive peak between 4 and 6 seconds and a negative post-undershoot at the 10.5 second time point. The responses were robust, with a peak between 0.2% and 0.4% for a single audiovisual word. The anterior STG preferred clear to noisy speech, while the posterior STG showed no preference or preferred noisy speech. To quantify these differences, linear mixed effects (LME) models were constructed.

Table 1 shows the results of the LME on the fMRI response amplitudes using an anatomically-defined border between anterior and posterior STG. There were three significant effects in the model. There was a small but significant effect of ROI location driven by a smaller overall response in the posterior STG (*p* = 0.01, effect magnitude of 0.07%). There were two larger effects: a main effect of auditory noise driven by a weaker overall response to noisy auditory stimuli (*p* = 10^−6^, magnitude 0.14%) and an interaction between auditory noise and ROI driven by a larger response to noisy auditory stimuli in posterior STG (*p* = 10^−4^, magnitude 0.16%).

**Table 1:** Linear Mixed-Effects Model of the Response Amplitude in STG Regions Defined 4 by Anatomical Boundary. Results of an LME model of the response amplitude in STG regions defined by an anatomical boundary. The fixed effects were the location of each region (Anterior *vs*. Posterior), the presence or absence of auditory noise (An) in the stimulus and the presence or absence of visual noise (Vn) in the stimulus. STG ROIs from right and left hemisphere across 6 subjects were included in the model as random factor. For each effect, the model estimate (in units of % signal change) for that factor is shown relative to baseline, the response in anterior STG ROI to clear audiovisual. The “Std. Error” column shows the standard error of the estimate. The degrees of freedom (“DF”), t-value and p-value derived from the model were calculated according to the Satterthwaite approximation, as provided by the *lmerTest* package (Kuznetsova et al 2015). The baseline is shown first, all other effects are ranked by absolute t-value. Significant effects are shown in bold.

| **Fixed Effects** | Estimate | Std. Error | DF | t-value | p-value |
| --- | --- | --- | --- | --- | --- |
| Baseline | 0.40 | 0.03 | 35.9 | 15.4 | $< 10^{-16}$ |
| **Auditory Noise (An)** | **-0.14** | **0.03** | **84** | **-5.4** | **$10^{-6}$** |
| **Posterior Location x An** | **0.16** | **0.04** | **84** | **4.2** | **$10^{-4}$** |
| **Posterior Location** | **-0.07** | **0.03** | **84** | **-2.6** | **0.01** |
| Visual Noise (Vn) | -0.03 | 0.03 | 84 | -1.0 | 0.31 |
| Posterior Location x Vn | -0.02 | 0.04 | 84 | -0.6 | 0.52 |
| Posterior Location x An x Vn | 0.01 | 0.05 | 84 | 0.2 | 0.83 |
| An x Vn | 0.00 | 0.04 | 84 | 0.0 | 0.97 |

Table 2 shows the results of an LME using a functionally-defined border between anterior and posterior STG. As with the LME on the anatomically-defined border, the largest effects were an interaction between auditory noise and location (*p* = 10^−4^, magnitude 0.21%) and a main effect of auditory noise (*p* = 10^−4^, magnitude −0.14%). These results are consistent with the LME on the anatomical border. One note of caution is that the LME using the functionally-defined border incorporated fMRI data, potentially biasing the model.

**Table 2:** Linear Mixed-Effects Model of the Response Amplitude in STG Regions Defined by Functional Boundary. Results of an LME model of the response amplitude in STG regions defined by the functionally-defined boundary between anterior and posterior STG.

| **Fixed Effects:** | Estimate | Std. Error | DF | t-value | p-value |
| --- | --- | --- | --- | --- | --- |
| Baseline | 0.37 | 0.03 | 38.32 | 10.8 | $10^{-13}$ |
| **Posterior Location x An** | **0.21** | **0.05** | **84** | **4.1** | **$10^{-4}$** |
| **Auditory Noise (An)** | **-0.14** | **0.04** | **84** | **-3.8** | **$10^{-4}$** |
| Posterior Location | -0.04 | 0.04 | 84 | -1.1 | 0.28 |
| Visual Noise (Vn) | -0.02 | 0.04 | 84 | -0.7 | 0.49 |
| Posterior Location x Vn | -0.02 | 0.05 | 84 | -0.4 | 0.66 |
| Posterior Location x An x Vn | 0.03 | 0.07 | 84 | 0.4 | 0.69 |
| An x Vn | -0.01 | 0.05 | 84 | -0.1 | 0.92 |

For comparison, Table 3 shows the results of a linear mixed-effects model on the ECOG responses, reprinted from Table 1 of (Ozker et al 2017). As with the LMEs on the fMRI data, the largest effects were a main effect of auditory noise (*p* = 10^−13^, magnitude - 110%) and an interaction between auditory noise and location (*p* = 10^−10^, magnitude +141%).

**Table 3:** Linear Mixed-Effects Model of the Response Amplitude in Anterior and Posterior ECOG electrodes. Results of an LME model of the ECOG response amplitude. The fixed effects were the location of each electrode (Anterior *vs*. Posterior), the presence or absence of auditory noise (An) in the stimulus and the presence or absence of visual noise (Vn) in the stimulus.

| Fixed effects: | Estimate | Std. Error | DF | t-value | p-value |
| --- | --- | --- | --- | --- | --- |
| **Baseline** | 183.1 | 24.8 | 33.7 | 7.4 | $10^{-8}$ |
| **Auditory noise (An)** | **-109.6** | **13.5** | **188** | **-8.1** | **$10^{-13}$** |
| **Posterior location x An** | **140.6** | **21.2** | **188** | **6.6** | **$10^{-10}$** |
| **Posterior location** | **-101** | **38.7** | **34.2** | **-2.6** | **0.01** |
| Visual noise (Vn) | 21.6 | 13.5 | 188 | 1.6 | 0.11 |
| An x Vn | -13.3 | 19.1 | 188 | -0.7 | 0.49 |
| Posterior location x Vn | -8.9 | 21.2 | 188 | -0.4 | 0.67 |
| Posterior location x An x Vn | 3.6 | 29.9 | 188 | 0.1 | 0.91 |

Direct comparison of the fMRI and ECOG responses required additional analyses. The first obstacle was the different time scale of the responses. Figure 4 shows the ECOG responses from the STG, reprinted from Figure 4 of (Ozker et al 2017). Increases in the high broadband signal began less than 100 ms after the onset of auditory speech, and peaked at about 200 ms after auditory speech onset. To convert the directly-recorded neural activity measured with ECOG to the indirect and much slower measure of neural activity provided by BOLD fMRI, the ECOG responses were convolved with a standard hemodynamic response function (Figure 4C) and downsampled from 1 ms resolution to a temporal resolution of 1.5 seconds, the repetition time (TR) of the fMRI data. This created a predicted fMRI response (based on the measured ECOG responses) on the same time scale and time base as the actual fMRI response. The second obstacle was the different amplitude scales of the responses. ECOG amplitude is measured in % change in the high-gamma broadband signal relative to pre-stimulus fixation baseline, while fMRI signal amplitude is measured in % intensity increase of the EPI images relative to fixation baseline. A separate scale factor was calculated for each condition in order to generate the best fit between the predicted and actual fMRI responses.

Figure 4D shows the predicted-from-ECOG responses and actual fMRI responses (based on the functional boundary between anterior and posterior STG). The shape of the responses was similar, as demonstrated by a high correlation coefficient between predicted and actual responses (anterior STG: 0.98 for auditory-clear, 0.96 for auditory-noisy; posterior STG: 0.97 for auditory-clear, 0.99 for auditory-noisy). The average across scale factor across conditions for the amplitude conversion was 612 ECOG% per BOLD%, meaning that a peak ECOG response of 612% was equivalent to a BOLD fMRI response of 1%. The scale factors were identical for auditory-clear and auditory-noisy conditions in posterior STG (476) but were markedly higher in anterior STG, especially for auditory-clear audiovisual words (909 for auditory-clear and 588 for auditory-noisy). This reflected the fact that in the ECOG data, the anterior STG response to clear speech was more than twice as large as the response to auditory-noisy speech (300% *vs*. 110%) while in fMRI, the anterior STG preferred clear speech to noisy speech but the difference was less pronounced (0.37% *vs*. 0.24%).

In ECOG and fMRI, we observed distinct patterns of responses to clear and noisy speech in anterior and posterior STG. A possible explanation for these results is that anterior STG is unisensory auditory cortex, rendering it susceptible to auditory noise added to speech, while posterior STG is multisensory auditory-visual cortex, allowing it to compensate for auditory noise using visual speech information. This explanation predicts that posterior STG should show stronger responses to visual speech than anterior STG. To test this explanation, we took advantage of the fact that the fMRI participants viewed standard block-design localizers containing sentences of unisensory visual, unisensory auditory and audiovisual speech. We measured the response to these localizers in STG ROIs defined using the functional boundary (Figure 5). This analysis was unbiased because the functional boundary was created using the auditory-clear and auditory-noisy data, completely independent of the localizers.

As predicted, the response to the unisensory visual speech presented in the localizer was significantly stronger in posterior STG than in anterior STG (posterior *vs*. anterior: 0.4% vs. 0.1%, p = 0.02 for visual speech). This could not be explained by an overall difference in responsiveness; in fact, for the other localizer stimuli, there was a trend towards weaker responses in posterior STG (posterior *vs*. anterior: 0.7% *vs*. 1.1%, *p* = 0.09 for unisensory auditory speech; 1.3% vs. 1.7%; *p* = 0.07 for audiovisual speech), consistent with the weaker responses in posterior STG to single audiovisual words observed in the main experiment (effect of posterior location in Table 1).

## Discussion

We measured neural activity in the human STG using two very different techniques: directly, using surface electrodes implanted in ECOG participants with epilepsy, or indirectly, using the BOLD response in fMRI participants were healthy controls. Both ECOG and fMRI participants viewed the same clear and noisy audiovisual speech stimuli and performed the same speech recognition task. Both techniques demonstrated a sharp functional boundary in the STG. On the anterior side of the boundary, cortex strongly preferred clear audiovisual speech to noisy audiovisual speech. On the posterior side of the boundary, cortex preferred noisy audiovisual speech or showed no preference. For both techniques, the boundary was located at a similar location in standard space (y = − 30 mm) and the transition between the two functional zones happened within 10 mm of anterior-to-posterior distance along the STG.

In both fMRI and ECOG patients, an anatomical boundary set at the most posterior point of Heschl’s gyrus provided a reasonable proxy for the functional boundary. This is important because unlike the fMRI or ECOG data needed to locate the functional boundary, the structural MRI scan needed to locate the anatomical boundary is easily obtainable (for instance, in the examination of patients with brain lesions). While primary visual and auditory cortex are easily localizable using anatomical landmarks, it has proven to be much more of a challenge to find landmarks for association areas (Weiner & Grill-Spector 2011, Witthoft et al 2014).

Multisensory integration provides a conceptual framework for understanding these results. When noisy auditory speech is presented, auditory information alone is insufficient for perception, and auditory-speech regions in anterior STG respond with diminished intensity. Visual speech information can compensate for noisy auditory speech (Bernstein et al 2004, Ross et al 2007, Sumby & Pollack 1954), but this requires recruitment of multisensory brain regions capable of combining the auditory and visual speech information to restore intelligibility. While both anterior and posterior STG responded to audiovisual speech, data from the fMRI localizer experiment showed that posterior STG responded more strongly to visual-only speech than anterior STG, supporting the idea that posterior STG is a multisensory area capable of combining auditory and visual speech.

The neural code in posterior STG is hinted at by a recent study that found that a region of posterior STG and STS (similar to the posterior STG region described in the present manuscript) preferred silent videos of faces making mouth movements to silent videos of faces making eye movements (Zhu & Beauchamp 2017). The same region responded strongly to unisensory auditory speech and preferred vocal to non-vocal sounds. Interestingly, as statistical thresholds were increased to select voxels with a greater preference for visual mouth movements, response to unisensory auditory speech increased, suggesting that at a single voxel level, small populations of neurons code for mouth movements and speech sounds, the two components of audiovisual speech (Bernstein et al 2011). This cross-modal correspondence in neural coding of multisensory cues is exactly as predicted by computational models of multisensory integration (Beck et al 2008, Magnotti & Beauchamp 2017).

There is a substantial body of evidence showing that posterior STS is a cortical hub for multisensory integration, multisensory, responding to both auditory and visual stimuli including faces and voices, letters and voices, and recordings and videos of objects (Beauchamp et al 2004, Calvert et al 2000, Foxe et al 2002, Miller & D’Esposito 2005, Reale et al 2007, van Atteveldt et al 2004). The finding that the adjacent cortex in posterior STG is also important for multisensory integration has several ramifications. In a transcranial magnetic stimulation (TMS) study, integration of auditory and visual speech (as indexed by the McGurk effect) was disrupted with TMS targeted at the posterior STS (Beauchamp et al 2010). The present results suggest that posterior STG may also have played a role in the observed disruption, and raise the possibility that electrical brain stimulation of STG in ECOG patients can increase our understanding of multisensory speech perception as it has for visual perception (Murphey et al 2008, Rangarajan & Parvizi 2015).

While there have been numerous previous fMRI studies of noisy and clear audiovisual speech, *e.g*. (Bishop & Miller 2009, Callan et al 2003, Lee & Noppeney 2011, McGettigan et al 2012, Sekiyama et al 2003, Stevenson & James 2009) none described a sharp boundary in the response patterns to clear and noisy speech within the STG. A likely explanation is that many of the previous studies use spatial filtering or blurring as a preprocessing step in their fMRI data analysis pipeline and reported only group average data, which introduces additional blurring due to inter-subject anatomical differences, especially for commonly-used volume-based templates. Combined, these two spatial blurring steps could easily eliminate sharp boundaries present in fMRI data. For instance, blurring eliminates the otherwise robust observation of functional specialization for different object categories in visual cortex (Tyler et al 2003). Another possible explanation for the failure of previous studies to observe the boundary is the common practice of reporting responses only at the location of activation peaks, rather than examining the entire extent of the activation. Anterior and posterior STG form a continuous region of active cortex, with the strongest activation in anterior STG. Therefore, only reporting responses from a single peak STG location (which would almost certainly fall in anterior STG) would camouflage the very different pattern of activity in posterior STG.

### Implications for ECOG and fMRI

While the primary goal of our study was not a comparison of the two methodologies, there was good correspondence between the actual fMRI signal and the fMRI signal predicted from our measure of ECOG amplitude, the broadband high-gamma response in the window from 70 - 110 Hz. This is consistent with mounting evidence that the high-frequency broadband signal in ECOG is a good match for the fMRI signal (reviewed in Ojemann et al 2013). Other ECOG measures, such as the narrowband gamma response (30 - 80 Hz) or the narrowband alpha response, may characterize neuronal synchrony rather than level of neuronal activity, and hence are poorly correlated with the BOLD signal (Hermes et al 2017).

A reassuring finding from the present study is that we observed similar patterns of responses between ECOG patients with epilepsy and healthy controls viewing the same stimuli and performing the same task. This provides data to partially mitigate persistent concerns that ECOG patients may have different brain organization than healthy controls, reducing the generalizability of the results of ECOG studies. One minor discrepancy between the ECOG and fMRI results was a larger relative amplitude for preferred stimuli in ECOG. For instance, anterior STG showed a nearly three-fold difference in the response amplitude to clear *vs*. noisy audiovisual speech (300% *vs*. 110%). The difference in fMRI was the same direction but much smaller (0.37% *vs*. 0.24%). We attribute this to the ability of ECOG electrodes to sample small populations of highly-selective neurons, while the BOLD fMRI response spatially sums over larger populations of neurons, mixing more and less selective signals. This same pattern has been observed in other studies comparing fMRI with ECOG. For instance, in a study of the fusiform face area, the BOLD signal evoked by faces was approximately double that evoked by non-face objects while the broadband high-gamma amplitude was triple or more for the same contrast (Parvizi et al 2012).

## Acknowledgments

We thank Johannes Rennig for assistance with fMRI data collection and Ping Sun, Daniel Yoshor and Bill Bosking for assistance with ECOG. Funding was provided by Veterans Administration Clinical Science Research and Development Merit Award Number 1I01CX000325-01A1 and NIH R01NS065395.

## References

Beauchamp MS, Lee KE, Argall BD, Martin A. 2004. Integration of auditory and visual information about objects in superior temporal sulcus. Neuron 41: 809–23

Beauchamp MS, Nath AR, Pasalar S. 2010. fMRI-Guided transcranial magnetic stimulation reveals that the superior temporal sulcus is a cortical locus of the McGurk effect. J Neurosci 30: 2414–7

Beck JM, Ma WJ, Kiani R, Hanks T, Churchland AK, et al. 2008. Probabilistic population codes for Bayesian decision making. Neuron 60: 1142–52

Belin P, Zatorre RJ, Lafaille P, Ahad P, Pike B. 2000. Voice-selective areas in human auditory cortex. Nature 403: 309–12

Bernstein LE, Auer ET, Takayanagi S. 2004. Auditory speech detection in noise enhanced by lipreading. Speech Communication 44: 5–18

Bernstein LE, Jiang J, Pantazis D, Lu ZL, Joshi A. 2011. Visual phonetic processing localized using speech and nonspeech face gestures in video and point-light displays. Human Brain Mapping 32: 1660–76

Bishop CW, Miller LM. 2009. A multisensory cortical network for understanding speech in noise. Journal of cognitive neuroscience 21: 1790–805

Callan DE, Jones JA, Munhall K, Callan AM, Kroos C, Vatikiotis-Bateson E. 2003. Neural processes underlying perceptual enhancement by visual speech gestures. Neuroreport 14: 2213–8

Calvert GA, Campbell R, Brammer MJ. 2000. Evidence from functional magnetic resonance imaging of crossmodal binding in the human heteromodal cortex. Current biology : CB 10: 649–57

Chang EF, Rieger JW, Johnson K, Berger MS, Barbaro NM, Knight RT. 2010. Categorical speech representation in human superior temporal gyrus. Nature neuroscience 13: 1428–32

Cheng K, Waggoner RA, Tanaka K. 2001. Human ocular dominance columns as revealed by high-field functional magnetic resonance imaging. Neuron 32: 359–74

Cox RW. 1996. AFNI: Software for analysis and visualization of functional magnetic resonance neuroimages. Comput Biomed Res 29: 162–73

Crosse MJ, Di Liberto GM, Lalor EC. 2016. Eye Can Hear Clearly Now: Inverse Effectiveness in Natural Audiovisual Speech Processing Relies on Long-Term Crossmodal Temporal Integration. J Neurosci 36: 9888–95

Dale AM, Greve DN, Burock MA. 5th International Conference on Functional Mapping of the Human Brain, Duesseldorf, Germany, 1999, 9.

Desikan RS, Segonne F, Fischl B, Quinn BT, Dickerson BC, et al. 2006. An automated labeling system for subdividing the human cerebral cortex on MRI scans into gyral based regions of interest. Neuroimage 31: 968–80

Destrieux C, Fischl B, Dale A, Halgren E. 2010. Automatic parcellation of human cortical gyri and sulci using standard anatomical nomenclature. NeuroImage 53: 1–15

Fischl B, Liu, A., Dale, A.M., 2001. Automated manifold surgery: constructing, the gaatcmo, human cerebral cortex. IEEE TMI 20 (1). 2001. Automated manifold surgery: constructing geometrically accurate and topologically correct models of the human cerebral cortex. IEEE TMI 20 (1), 70–80. IEEE Transactions in Medical Imaging 20: 70–80

Foxe JJ, Wylie GR, Martinez A, Schroeder CE, Javitt DC, et al. 2002. Auditory-somatosensory multisensory processing in auditory association cortex: an fMRI study. Journal of neurophysiology 88: 540–3

Grill-Spector K, Malach R. 2004. The human visual cortex. Annu Rev Neurosci 27: 649–77

Hermes D, Nguyen M, Winawer J. 2017. Neuronal synchrony and the relation between the blood-oxygen-level dependent response and the local field potential. PLoS Biol 15: e2001461

Holmes CJ, Hoge R, Collins L, Woods R, Toga AW, Evans AC. 1998. Enhancement of MR images using registration for signal averaging. J Comput Assist Tomogr 22: 324–33

Janszky J, Jokeit H, Heinemann D, Schulz R, Woermann FG, Ebner A. 2003. Epileptic activity influences the speech organization in medial temporal lobe epilepsy. Brain 126: 2043–51

Kramer MA, Cash SS. 2012. Epilepsy as a disorder of cortical network organization. Neuroscientist 18: 360–72

Kuznetsova A, Brockhoff PB, Christensen RHB. 2015. Package ‘lmerTest’. R package version: 2.0–29

Leaver AM, Rauschecker JP. 2016. Functional Topography of Human Auditory Cortex. J Neurosci 36: 1416–28

Lee H, Noppeney U. 2011. Physical and perceptual factors shape the neural mechanisms that integrate audiovisual signals in speech comprehension. The Journal of neuroscience : the official journal of the Society for Neuroscience 31: 11338–50

Lindquist MA, Meng Loh J, Atlas LY, Wager TD. 2009. Modeling the hemodynamic response function in fMRI: efficiency, bias and mis-modeling. Neuroimage 45: S187–98

Magnotti JF, Beauchamp MS. 2017. A Causal Inference Model Explains Perception of the McGurk Effect and Other Incongruent Audiovisual Speech. PLoS Comput Biol 13: e1005229

McGettigan C, Faulkner A, Altarelli I, Obleser J, Baverstock H, Scott SK. 2012. Speech comprehension aided by multiple modalities: behavioural and neural interactions. Neuropsychologia 50: 762–76

Mesgarani N, Cheung C, Johnson K, Chang EF. 2014. Phonetic feature encoding in human superior temporal gyrus. Science 343: 1006–10

Miller LM, D’Esposito M. 2005. Perceptual fusion and stimulus coincidence in the cross-modal integration of speech. The Journal of neuroscience : the official journal of the Society for Neuroscience 25: 5884–93

Moerel M, De Martino F, Formisano E. 2014. An anatomical and functional topography of human auditory cortical areas. Front Neurosci 8: 225

Murphey DK, Yoshor D, Beauchamp MS. 2008. Perception matches selectivity in the human anterior color center. Current biology : CB 18: 216–20

Nath AR, Beauchamp MS. 2012. A neural basis for interindividual differences in the McGurk effect, a multisensory speech illusion. NeuroImage 59: 781–7

Nath AR, Fava EE, Beauchamp MS. 2011. Neural correlates of interindividual differences in children’s audiovisual speech perception. J Neurosci 31: 13963–71

Ojemann GA, Ojemann J, Ramsey NF. 2013. Relation between functional magnetic resonance imaging (fMRI) and single neuron, local field potential (LFP) and electrocorticography (ECoG) activity in human cortex. Front Hum Neurosci 7: 34

Ozker M, Schepers IM, Magnotti JF, Yoshor D, Beauchamp MS. 2017. A Double Dissociation between Anterior and Posterior Superior Temporal Gyrus for Processing Audiovisual Speech Demonstrated by Electrocorticography. J Cogn Neurosci 29: 1044–60

Parvizi J, Jacques C, Foster BL, Witthoft N, Rangarajan V, et al. 2012. Electrical stimulation of human fusiform face-selective regions distorts face perception. J Neurosci 32: 14915–20

Rangarajan V, Parvizi J. 2015. Functional asymmetry between the left and right human fusiform gyrus explored through electrical brain stimulation. Neuropsychologia

Ray S, Maunsell JH. 2011. Different origins of gamma rhythm and high-gamma activity in macaque visual cortex. PLoS biology 9: e1000610

Reale RA, Calvert GA, Thesen T, Jenison RL, Kawasaki H, et al. 2007. Auditory-visual processing represented in the human superior temporal gyrus. Neuroscience 145: 162–84

Ross LA, Saint-Amour D, Leavitt VM, Javitt DC, Foxe JJ. 2007. Do you see what I am saying? Exploring visual enhancement of speech comprehension in noisy environments. Cerebral cortex (New York, N.Y. : 1991) 17: 1147–53

Salmelin R. 2007. Clinical neurophysiology of language: the MEG approach. Clin Neurophysiol 118: 237–54

Sekiyama K, Kanno I, Miura S, Sugita Y. 2003. Auditory-visual speech perception examined by fMRI and PET. Neuroscience research 47: 277–87

Shahin AJ, Kerlin JR, Bhat J, Miller LM. 2012. Neural restoration of degraded audiovisual speech. Neuroimage 60: 530–8

Sohoglu E, Davis MH. 2016. Perceptual learning of degraded speech by minimizing prediction error. Proc Natl Acad Sci U S A 113: E1747–56

Stevenson RA, James TW. 2009. Audiovisual integration in human superior temporal sulcus: Inverse effectiveness and the neural processing of speech and object recognition. Neuroimage 44: 1210–23

Sumby WH, Pollack I. 1954. Visual contribution to speech intelligibility in noise. The journal of the acoustical society of america 26: 212–15

Tang C, Hamilton LS, Chang EF. 2017. Intonational speech prosody encoding in the human auditory cortex. Science 357: 797–801

Tyler LK, Bright P, Dick E, Tavares P, Pilgrim L, et al. 2003. Do semantic categories activate distinct cortical regions? Evidence for a distributed neural semantic system. Cogn Neuropsychol 20: 541–59

van Atteveldt N, Formisano E, Goebel R, Blomert L. 2004. Integration of letters and speech sounds in the human brain. Neuron 43: 271–82

Weiner KS, Grill-Spector K. 2011. Not one extrastriate body area: using anatomical landmarks, hMT+, and visual field maps to parcellate limb-selective activations in human lateral occipitotemporal cortex. Neuroimage 56: 2183–99

Witthoft N, Nguyen ML, Golarai G, LaRocque KF, Liberman A, et al. 2014. Where is human V4? Predicting the location of hV4 and VO1 from cortical folding. Cereb Cortex 24: 2401–8

Zhu LL, Beauchamp MS. 2017. Mouth and Voice: A Relationship between Visual and Auditory Preference in the Human Superior Temporal Sulcus. J Neurosci 37: 2697–708

